# Genomic Reconstruction and Dietary Response Assessment of Three *Clostridium leptum*-related Bacteria Isolated from Fecal Samples of Singapore Subjects

**DOI:** 10.1101/2024.09.13.612987

**Authors:** Mi Ae Park, Sharifah Nora Ahmad Almunawar, Rachel Rui Xia Lim, Sumanto Haldar, Christiani Jeyakumar Henry, Oleg Vladimirovich Moskvin

## Abstract

*Clostridium leptum*, a key player in gut butyrate production, has a profound impact on various facets of intestinal health. A recent clinical trial highlighted a significant increase in the relative abundance of this species in response to dietary interventions using beneficial oils. We isolated microbial strains corresponding to “*Clostridium leptum*” (at the 16S rRNA gene similarity level) and sequenced their genomes. All three genomes were successfully reconstructed, maintaining the chromosome as a single contig. Subsequent genome-wide analysis unveiled the phylogenetic diversity of the isolates, including the discovery of a new species - *Gallacutalibacter singaporense*. Based on the reconstructed metabolic model, we predicted the growth condition patterns of this new species and confirmed the predictions *in vitro*. Leveraging the assembled genomes, we dissected the components of the strong dietary intervention response signal previously ascribed to “*C. leptum*” and revealed distinct individual dynamics of all three bacteria in the clinical trial context. The transitional behavior of the novel species, in particular, revealed intriguing patterns, blazing the path to uncovering previously unrecognized interactions along the diet – gut microbiome – human health axis.

## 2. Introduction

*Clostridium leptum* plays a pivotal role in butyrate production within the gut, which in turn has been linked to numerous benefits for intestinal health and beyond (Louis and Flint 2009, Canani, Costanzo et al. 2011). The range of its beneficial effects on human health also extends to immunity modulation, such as the reduction of airway inflammation in asthma via stimulation of regulatory B cells (Huang, Zhang et al. 2022). *C*.*leptum* abundance decreases in inflammatory bowel disease (Cardoneanu, Mihai et al. 2021), oncological cases (Gui, Li et al. 2020), and COVID-19 patients (Tang, Gu et al. 2020). Together with other beneficial bacteria in the gut, *C*.*leptum* is induced by a *Lactobacillus* probiotic intervention (Mei, Zheng et al. 2021), and also a Mediterranean diet; the latter is attributed to a combined effect of polyunsaturated fatty acids and dietary fiber (Barber, Kabisch et al. 2023). In a recent clinical trial, we investigated the microbiome effects of a dietary intervention using beneficial oils (Lim, Park et al. 2022). One significant finding was a pronounced increase in the relative abundance of this species, mirroring the dietary intervention dynamics. The present paper delves into our follow-up research leveraging a combination of culturomics and *in silico* modeling. We successfully isolated three distinct microbial strains corresponding to “*Clostridium leptum*” at low-resolution (16S rRNA gene) level, reconstructed their genomes and demonstrated unique clinical association patterns and wider than expected phylogenetic diversity, spanning different species and even genera. This achievement contributes to understanding this taxonomic branch and its role within the human gut.

## 3. Methods

### 3.1. Bacteria isolation

Bacterial isolation involved the extraction of viable bacteria from frozen fecal samples that had been initially collected as part of the gut microbiome sub-project (Lim, Park et al. 2022), an extension of the previously documented dietary intervention clinical trial (Haldar, Wong et al. 2020). Sample selection involved (a) metagenomic-based pre-selection of high-responding subjects and (b) use of final time point samples for pre-selected subjects to enrich for strains most responsive to the intervention. To initiate the isolation process, 20 mg of frozen material was carefully resuspended under anaerobic conditions in 500 µl of the growth medium (described below). Subsequently, the resuspended material was plated onto Petri dishes containing an agar-amended version of the same medium. The formulation of the growth medium was derived from ATCC2751 maltose medium, with specific adjustments: firstly, the concentration of L-Cysteine was doubled (1 g/l), and secondly, an additional 0.3 g/l of Pyruvate was incorporated to enhance the medium’s efficacy.

Individual colonies were cultivated under anaerobic conditions in 10 ml of the above medium. This culture served a dual purpose: glycerol stocks for biobanking and DNA isolation for further analysis.

### 3.2. DNA isolation and sequencing

We extracted DNA from bacterial cultures using the QIAGEN Power Fecal kit (Qiagen, USA), following the manufacturer’s guidelines and the following custom variations: 1) for the metagenomic sequencing of the clinical samples, the detailed procedure was described earlier (Lim, Park et al. 2022); 2) for genome reconstruction of individual strains, considering the downstream nanopore sequencing involved, DNA isolation conditions were optimized for maximum retention of the long DNA fragments. The DNA was extracted from a broth bacterial culture derived from a pure single colony, and the homogenization step upstream of the QIAamp PowerFecal kit was conducted using a Bead Beater (MP Biomedicals, USA) at 6 m/s for 15 seconds; 3) for the extensive screening of colonies via 16S rRNA Gene PCR, we used fast DNA purification from a bacterial culture derived from a pure single colony, using the InstaGene matrix (Bio-Rad, USA) according to the manufacturer’s instructions.

The 16S rRNA gene PCR was conducted targeting the 16S rRNA gene region from position 27 to 1429, using the following primers: 27F (5’-AGAGTTTGATCCTGGCTCAG-3’) and 1429R (5’-ACGGCTACCTTGTTACGACTT-3’). PCR conditions included an initial denaturation at 95°C for 2 minutes, followed by 32 cycles of 95°C for 20 seconds, 55°C for 20 seconds, and 72°C for 1 minute, and a final extension at 72°C for 7 minutes.

Short-read sequencing of both the clinical samples and individual isolates was performed using the Illumina HiSeq2000 platform. We constructed the sequencing library using 1.5 µg of purified DNA, with specifications of 2 x 150bp reads and a 300 bp insert.

For long-read sequencing, we utilized the MinION device from Oxford Nanopore. The sequencing library was prepared as per the manufacturer’s instructions for the LSK-110 sequencing kit.

### 3.2. Genome assembly and analysis

Prior to assembly, we subjected the sequencing reads to a quality filtering process. For Illumina reads, we employed *Trimmomatic* (Bolger, Lohse et al. 2014) version 0.33, applying stringent parameters: a minimum average Q-score of 32 within a sliding window of 3 and a post-trim read length of at least 135 bp. For Nanopore long reads, *Nanofilt* (De Coster, D’Hert et al. 2018) was used to retain reads with a minimum quality of Q10 and a length threshold of 18 Kilobases. Though the length filtering may seem stringent, it aligns with our internal assessments on the influence of this parameter on genome assembly quality. The assembly was performed with Unicycler (Wick, Judd et al. 2017) version 0.4.9. In all the 3 cases discussed in this paper, the chromosome was recovered as a single contig. The assembled genomes were annotated with *Prokka* (Seemann 2014). Automated whole-genome metabolic model building was performed using *CarveMe* (Machado, Andrejev et al. 2018). Application of the metabolic model for growth conditions prediction was accomplished with COBRA toolbox (Heirendt, Arreckx et al. 2019). Phylogenetic assignments based on the entire genome information were performed using GTDB-Tk (Chaumeil, Mussig et al. 2022) version 2.4.0 (April 2024).

### 3.3. Estimating the relative contribution of strains in metagenomic signatures

The first-pass phylogenetic screening using 16S rRNA gene assigned all the three bacterial strains we isolated from the clinical study subjects to a single record – “*Clostridium leptum*”. The same is true for marker-based phylogenetic assignments of the metagenomic reads generated in the clinical trial. To assess the origin of the metagenomic reads by leveraging the reconstructed genomes, we took a closer look at the metagenomic sequencing data from the clinical study by adapting the RNA-Seq read mapping algorithm, RSEM (Li and Dewey 2011). While RSEM was initially designed to tackle issues arising from multiple mapping positions of the same sequencing read in repetitive genomic regions, we repurposed its probabilistic read mapping capabilities to accurately assign reads to closely related reference genomes. We fed the algorithm with the three reconstructed reference genomes. It then processed every read from the 378 FASTQ files (Lim, Park et al. 2022) to estimate the contribution of each of the three bacteria to the pool of metagenomic sequences in each sample. This procedure enabled us to ascertain which genome each read originated from. As an output from RSEM, we obtained a Posterior Mean Estimate of counts for each of the 1,134 sample-genome pairs. We then normalized these counts against the total number of sequencing reads in the corresponding samples. This process gave us the relative abundance of the three strains in every sample.

Differential abundance of the tree strains relevant to the clinical trial (temporal changes and differences between the study groups) were assessed in a multivariate analysis setting using *Maaslin2* (Morgan, Tickle et al. 2012) version 1.3.2, similar to global analysis of the entire metagenomic data in the original clinical trial. The parameters of interest (Group and Time) were set as target variables, and subject ID as a random effect.

## 4. Results and Discussion

### 4.1. Comparison of reconstructed genomes of the C.leptum related strains

Table 1A shows the overview of 3 reconstructed bacterial genomes showing “*Clostridium leptum* DSM753” as the closest 16S rRNA-based taxonomic entity, as well as the reference *C.leptum* DSM753 genome available from NCBI. It is worth noting that unlike fragmentary reconstruction of the currently available reference genome (22 contigs), our reconstructions yielded chromosomes as a single contig in all the 3 cases. This result can be attributed to a combination of a) hybrid assembly that leverages both short and long reads and b) large sequencing depth (around 3 Gbp per sample of both Illumina and Nanopore sequences) that allowed us to apply stringent quality filtering to both of the sequencing read types before the assembly. Among the 3 isolated strains, one matches 100% to the known *C.leptum* strain (DSM753) at the 16S rRNA gene level, while two others differ significantly even at the genome size level. The bacterium with the larger genome and notoriously slow growth rate (14 days were required for the colony formation, and the same time required for the growth in the liquid culture) had only 95% identity of its 16S rRNA gene, significant divergence in the gene content (Table 1B) and hence can be, even conservatively, attributed to a new species or, rightfully, to a new genus. We leveraged the reconstructed genomes to obtain more accurate phylogenetic placement via GTDB-Tk pipeline. The results are presented in the right column of Table 1A. While species assignments for the 2 fast-growing isolates was unequivocal (*Clostridium leptum* and *Clostridium sp902490005)*, the slow-growing isolate with larger genome and the lowest identity level for the 16S rRNA gene, was confidently assigned to *Gallacutalibacter* genus, representing a novel species within that genus (RED value 0.844). We called it *Gallacutalibacter singaporense*. The smaller genome size of *C. sp902490005* was confirmed by parallel genome reconstruction of two separate clones, including parallel Illumina and Nanopore sequencing and the entire quality filtering and genome assembly pipelines (Table 1A). This smaller-genome strain is designated *C. sp902490005* SG001. The entire story underlines the limitations of the 16S rRNA-based phylogeny placements, with quite an extreme case of this study when 3 isolates that have the identical top match (*C.leptum* DSM 753), in fact, belong to 3 different species and moreover, 2 different genera.

**Table 1.** **A**, 16S rRNA gene identity levels, genome assembly statistics and genome-wide phylogenetic placement results (GTDB-tk) for the 3 isolated strains and the available NCBI reference. For the GTDB-tk, the associated values of ANI (Average Nucleotide Identity), AF (Alignment Fraction) and RED (Redundancy Estimation for the Detection of Novel Taxa) are reported. Multiple Sequence Alignment scores (MSA, not shown) were above 95% for all the 3 reconstructed genomes. For the strains isolated in this study, the “Source” barcodes were defined as a concatenation of the study ID (“BOB”), the subject number (“NNN”) and the clone number for that subject (“CNNN”), where “N” is a digit. **B**, Statistics of protein-level genome annotation results.

| Source | Top hit (16S)<br>/ identity<br>level | Contigs | Assembly<br>size, Kbp | Growth<br>in the<br>culture | GTDB-tk placement / ANI / AF / RED /<br>proposed name |
| --- | --- | --- | --- | --- | --- |
| NCBI Reference<br>1263068.PRJEB802 | *Clostridium*<br>*leptum* DSM<br>753 / n/a | 22 | 3,270 | fast | n/a |
| BOB Study<br>(BOB030C018) | *Clostridium*<br>*leptum* DSM<br>753 / 100% | 1 | 3,275 | fast | *Clostridium leptum* / 98.5 / 0.80 / n/a<br>*Clostridium leptum* |
| BOB Study<br>(BOB030C027<br>BOB030C028) | *Clostridium*<br>*leptum* DSM<br>753 / 96% | 1 | 3,093<br>3,093 | fast | *C. sp902490005* / 98.2 / 0.91 / n/a<br>*Clostridium sp902490005* SG001 |
| BOB Study<br>(BOB136C69) | *Clostridium*<br>*leptum* DSM<br>753 / 95% | 2 | 3,565 | slow | *Gallacutalibacter* / n/a / n/a / 0.844<br>*Gallacutalibacter singaporense* |

**B.**
| Singapore isolate | ORFs<br>total | Annotated<br>ORFs | Unique gene<br>names | Loss<br>compared to<br>DSM753 | Gain compared<br>to DSM753 |
| --- | --- | --- | --- | --- | --- |
| *Clostridium leptum* DSM 753 | 2,944 | 1,364 | 1,014 | - | - |
| *Clostridium leptum* SG 001 | 2,730 | 1,418 | 1,018 | 106 | 110 |
| *Clostridium singaporense* | 2,868 | 1,302 | 1,012 | 235 | 233 |

### 4.2. Peculiarities of G.singaporense: in vitro growth

Paradoxically at the first glance, as already mentioned, the larger-genome and gene-rich *C.singaporense* demonstrated an extremely slow growth rate, showing the possible existence of metabolic impairment and high dependence of this strain on the microbial community supplying intermediate metabolites. Essentially, this bacterium must represent the nearly “unculturable”, highly specialized part of the gut microbiome, and its isolation in the first place was possible by a combination of highly permissive growth medium (with addition of pyruvate) and special attention paid to extremely slow growing colonies. At the level of gene content, we see the reason for that (Table 2): the central carbon metabolism is impaired by absence of both transketolase / transaldolase and all subunits of the dihydroxyacetone kinase. At the same time, compared to *C.leptum* DSM753, *G.singaporense* has a richer potential to metabolize amino acids, as well as withstand oxidative stress and heavy metal contamination (Table 2). And also, unlike DSM 753, it possesses more genes encoding Glycerol-3-P related enzymes, including both its transporter and another form of Glycerol-3-P dehydrogenase. The phenomenon of partial compensation of the absence of dihydroxyacetone kinase by an active glycerol kinase was described in the literature long ago (Jin, Forage et al. 1982).

**Table 2.** Comparison of critical differences in enzyme-encoded genes between *G.singaporense* and *C.leptum* DSM753. +/− stand for the presence or absence of the respective gene in the annotated genome.

| Enzyme-encoding gene | *Gallacutalibacter*<br>*singaporensis* | *Clostridium*<br>*leptum* DSM 753 |
| --- | --- | --- |
| Dihydroxyacetone kinase (all subunits) | - | + |
| Transaldolase | - | + |
| Transketolase | - | + |
| Lysine decarboxylase | + | - |
| Methionine transporter | + | - |
| Threonine dehydrogenase | + | - |
| Alkyl hydroperoxide reductase | + | - |
| Superoxide dismutase (Cu-Zn) | + | - |
| Superoxide dismutase (Mn) | + | - |
| Mercuric reductase | + | - |
| Zinc Sigma-54 two component system | + | - |
| Multidrug and toxin extrusion (MATE) pump<br>YdhE | + | - |
| Glycerol kinase | + | + |
| Glycerol-3-P dehydrogenase [NADP+] EC1.1.1.94 | + | + |
| Glycerol-3-P ABC transporter (both subunits) | + | - |
| Glycerol-3-P dehydrogenase EC1.1.5.3 | + | - |
| NADH peroxidase | + | - |

Going from consideration of individual genes to a more holistic view on the strain’s metabolic potential, we reconstructed the whole-genome metabolic model of *G.singaporense* and ran several versions of prediction of the growth requirements using the generated model within the COBRA toolbox. Several growth media predictions, obtained with different settings for minimizing the number of components and relative growth rate requirement, had common themes: Arabinose, Oxygen and Nitrate (Fig. 1). The predicted ability to use alternative terminal electron acceptors and tolerance to oxygen were intriguing, and we checked the model’s recommendations *in vitro*.

**Figure 1.**
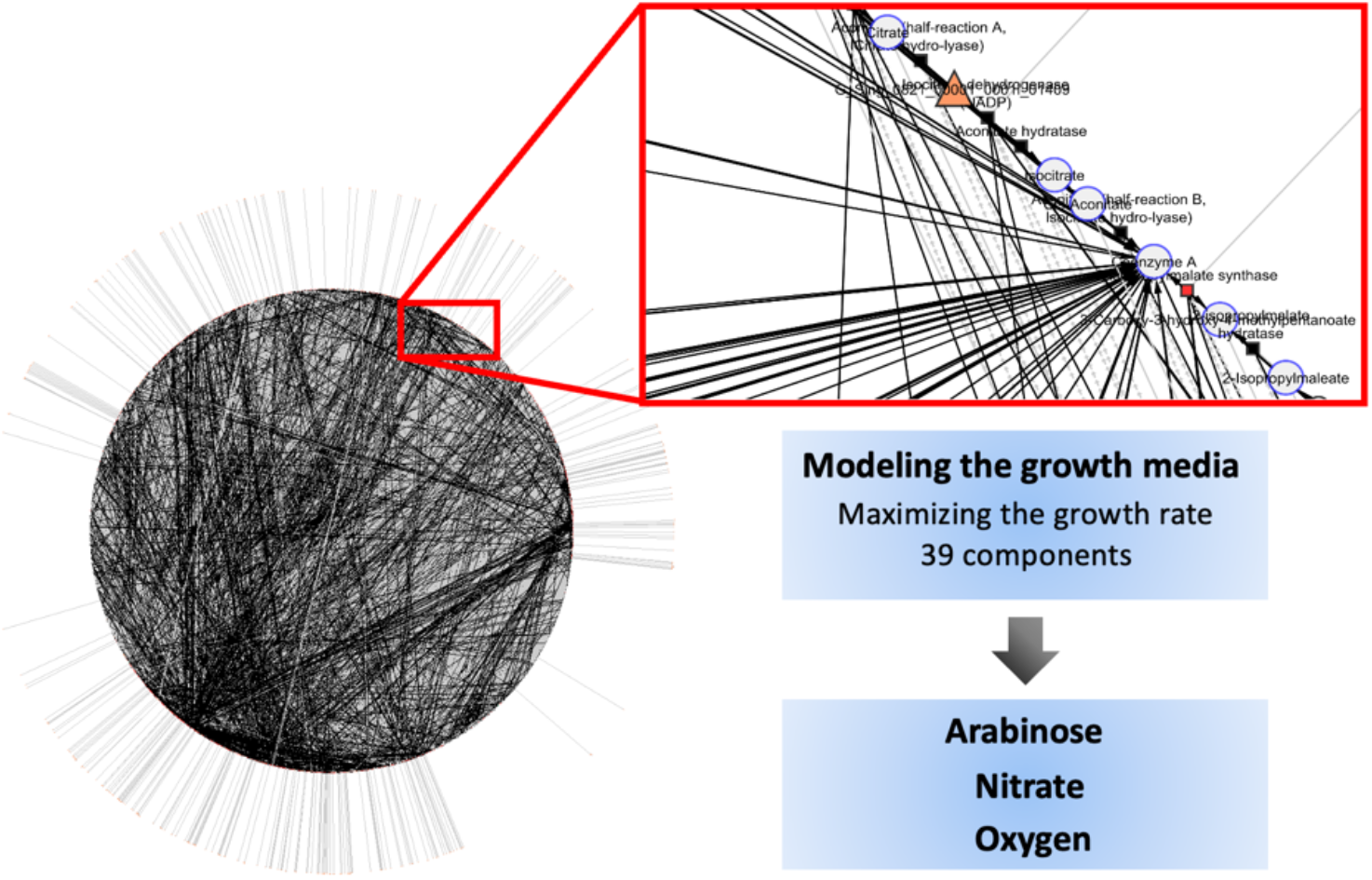
The whole-genome metabolic model of *G.singaporense* (presented as a network of enzymes and metabolites) and the main growth condition themes suggested by the model when predicting the minimal growth medium.

Fig. 2 shows growth curves of *G.singaporense* recorded for 250 hrs incorporating the *in silico* recommendations into *in vitro* growth conditions. A notable finding, aligned with the model’s prediction, was the more than five-fold enhancement in growth upon replacing Maltose with Arabinose. This observation was consistent in both anaerobic (Fig. 2A) and aerobic (Fig. 2C) environments. Introducing nitrate as an alternative electron sink led to interesting patterns: in the presence of both nitrate and oxygen, initial growth rates on Maltose and Arabinose were nearly equivalent. However, over time, there was a preferential inclination towards Arabinose, with cultures using Maltose showing decay (Fig. 2D). In strict anaerobic conditions with only nitrate, an unexpected surge in bacterial growth—nearly five times compared to other conditions—was observed when Maltose was present (Fig. 2B). The inherent dynamics causing the latter phenomena remain unexplained by the metabolic model. It’s imperative to realize that this model would benefit from comprehensive manual adjustments and the integration of empirically determined kinetic parameters. Such developments (representing a separate project) would enhance the model’s capacity to describe more intricate responses observed *in vitro*.

**Figure 2.**
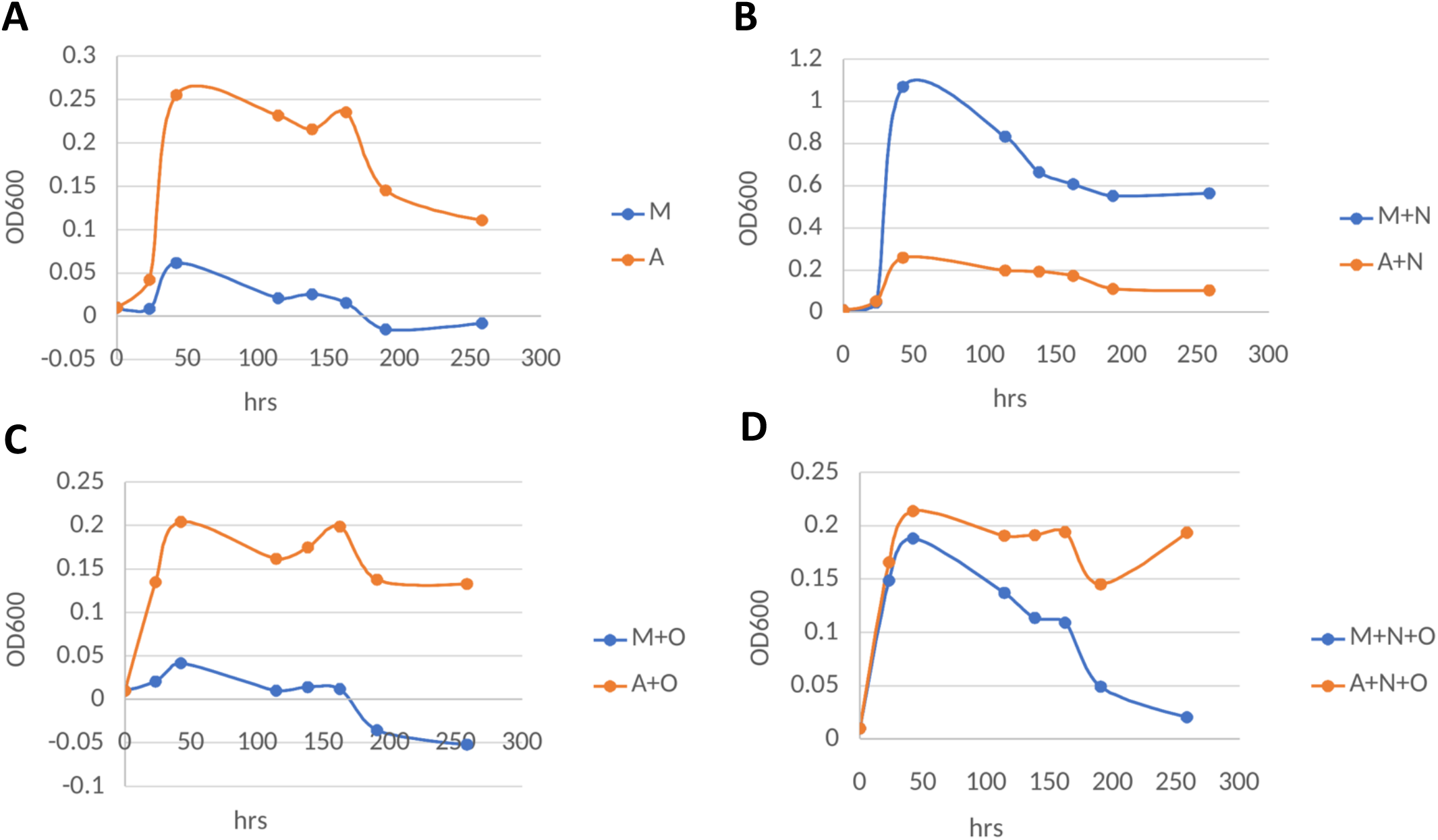
Growth curves of in vitro cultures of G.*singaporense* grown anaerobically **(A, B)** or aerobically **(C, D)**, in the absence **(A, C)** or presence **(B, D)** of nitrate in the medium. In-panel abbreviations: M, maltose; A, arabinose; O, oxygen; N, nitrate.

### 4.3. Estimation of the relative contributions of the 3 bacteria to “C.leptum” metagenomic signal

Following the methodology outlined earlier, we utilized the multimapping-aware read mapping algorithm to differentiate among related genomes and probabilistically designate each sequencing read to its genuine genome of origin, also bypassing the limitations of the absence of a newly reconstructed genomes in any database. A subsequent correlation test between the abundances of our three target bacteria and the clinical trial parameters unearthed a pattern reminiscent of previously published data: there was no correlation with the “Group” variable, yet there were notable associations with the “Time” variable for two of the three *Clostridium leptum*-related strains, as depicted in Figure 3. A more granular perspective detailing sample-centric relative alterations between pairs of pertinent strains at Day 14 and Day 56 can be found in Figure 4. Three significant observations arose: 1) The dominant impact of the “*C.leptum*” induction can be attributed to the widely recognized strain DSM753, 2) The new strain SG001 doesn’t exhibit a systematic effect across the entire study population, 3) Although *G.singaporense* is the least abundant among the trio, its trajectory during the trial is notable. It experienced suppression on Day 14, followed by induction by Day 56, ultimately presenting an increasing trend as the study concluded. This trend was corroborated by a formally positive association coefficient as per Maaslin2 mixed model analysis. The most prominent response to the dietary intervention with the beneficial oils still belongs to the well-established *Clostridium leptum* species (Figure 4). Notably, for a small subset of the subjects that demonstrated the opposite response to the dietary intervention, compared to the main trend, i.e., decrease in abundance, similar depletion in abundance was observed in both of the other strains (Fig4, lower left quadrants), which is especially prominent at Day 56 (Panels B and D), where for the 5 subjects with the strongest negative trend for *C.leptum*, abundance changes for the other two species were also negative, underlying community relationships within this taxonomic group, fine details of which deserve a focused study.

**Figure 3.**
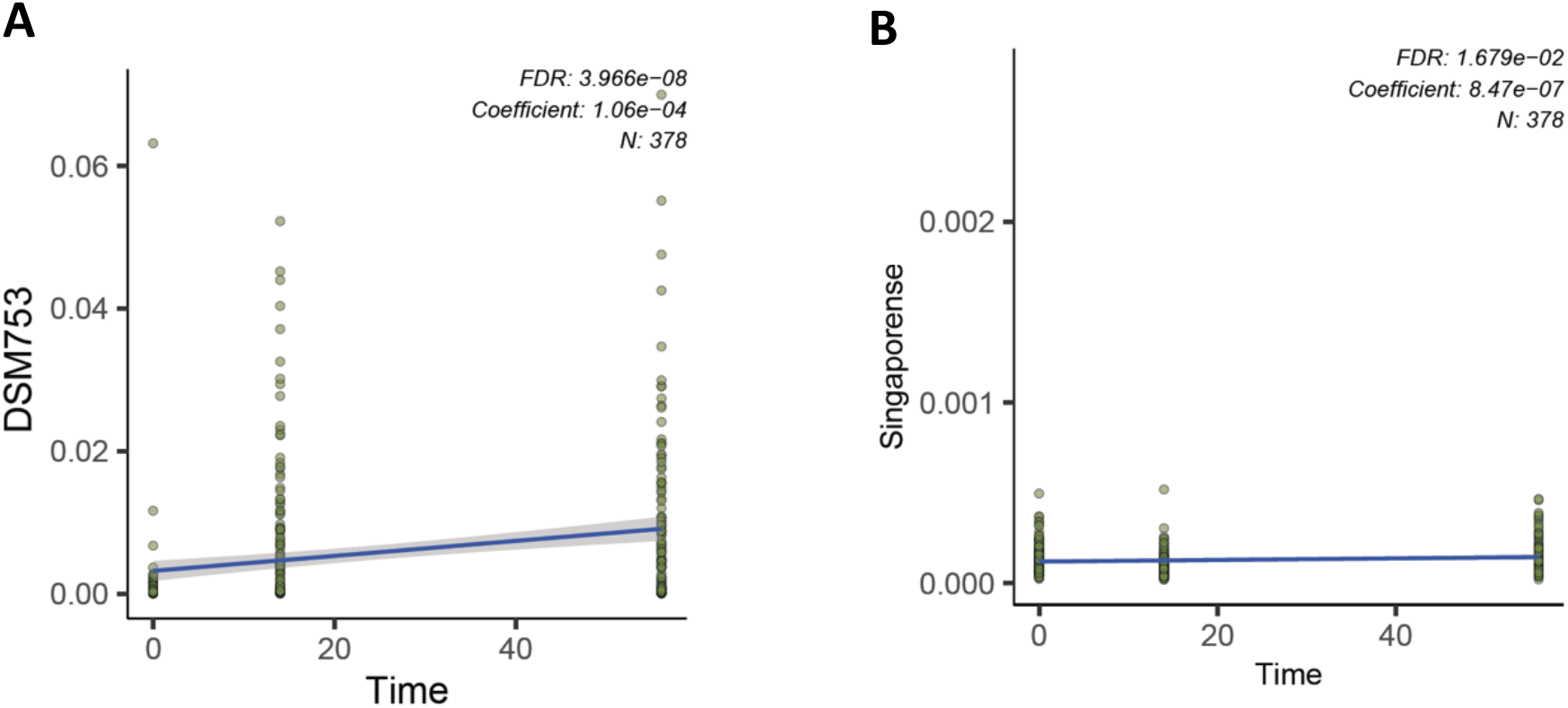
Association results between the time of the dietary intervention and the relative abundance of *C.leptum* DSM753 **(A)** and *G.singaporense* **(B)**, according to the mixed model implemented in *Maaslin2*.

**Figure 4.**
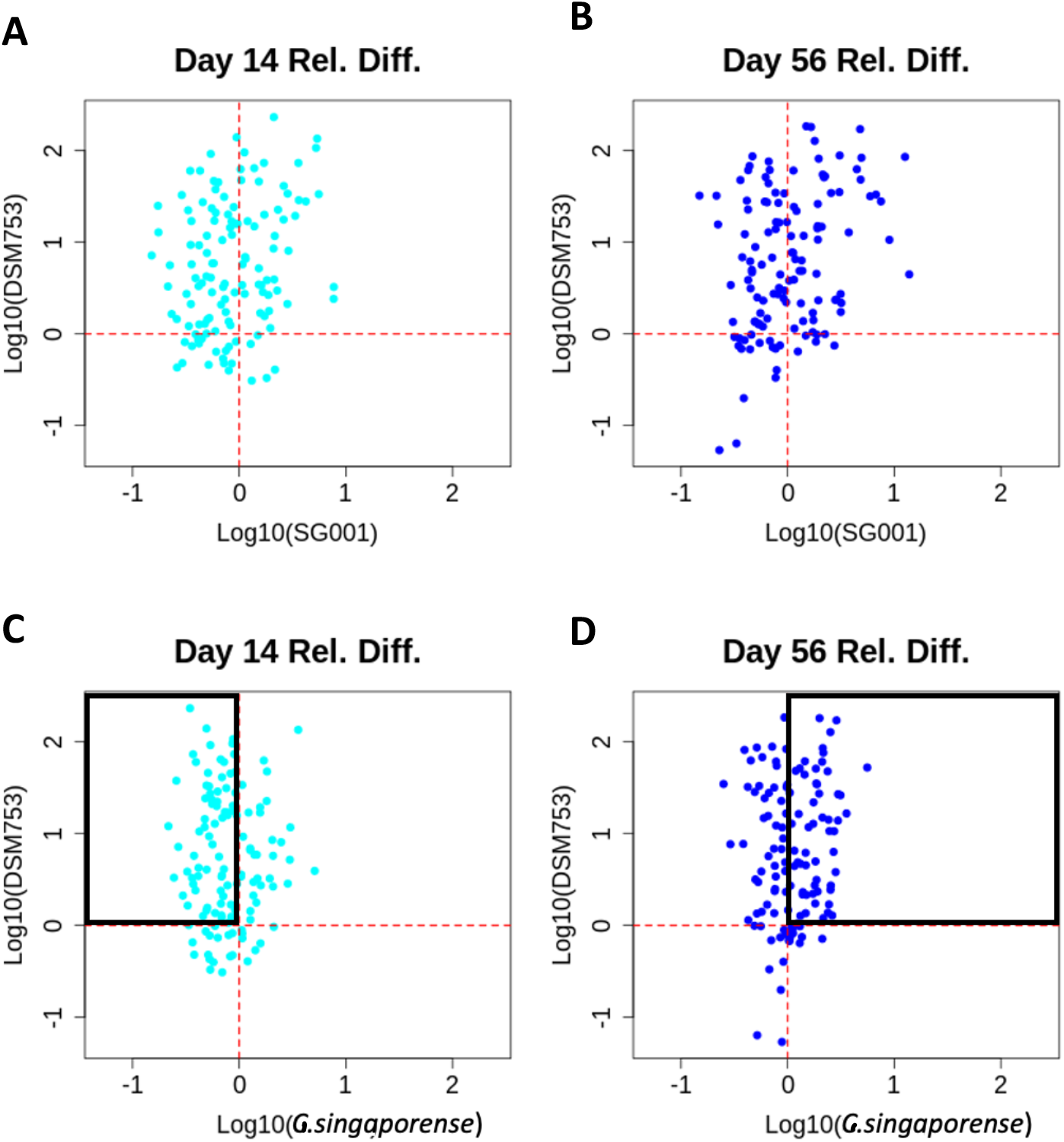
Differential abundance of *C.leptum* DSM 753 vs. either *C.sp902490005* SG001 **(A, B)** or *G.singaporense* **(C, D)** at Day 14 **(A, C)** or Day 56 **(B, D)**, relative to the pre-intervention state (Day 0). Rectangles highlight the transitional response of *G.singaporense* in the larger subset of the subjects where *C.leptum* DSM753 demonstrated unidirectional and early-saturated response.

## 5. Conclusion

We successfully isolated three unique bacterial strains, with one identified as a novel species, all close in taxonomy space to *Clostridium leptum*, being indistinguishable with low-resolution 16S rRNA-based taxonomy mapping. We assembled their high-quality reference genomes, with chromosome in a single contig, and analyzed the key peculiarities of gene content and the preliminary reconstructed metabolic model to predict *in vitro* growth behavior of the new species – *Gallacutalibacter singaporense*.

The distinct patterns of associations with the clinical trial dynamics, especially the notable transitional behavior of *G.singaporense*, could hint at previously undiscovered microbial or host-microbe interactions. These dynamics emphasize the intricate relationships between these dietary responsive bacteria, which, before our research, were collectively and indistinctly labeled under the “*Clostridium leptum*” moniker, according to lower-resolution 16S metabarcoding data. Delving deeper into these community relationships may have important implications for clinical outcomes, considering the unusually dramatic increase in the relative abundance of *C.leptum* in response to the beneficial oils intervention and the respective increase in butyrate-producing genome load of the community (Lim, Park et al. 2022).

## 7. Data Availability

The data that support the findings of this study, including both the metagenomic sequences and the reference genomes for the bacterial isolates, are openly available in NCBI Sequence Read Archive, BioProject PRJNA728374.

## 8. Acknowledgements

We thank Prof. Chua Nam Hai for his expert guidance throughout the project and comments on the manuscript. The computational work for this article was partially performed on resources of the National Supercomputing Centre, Singapore (https://www.nscc.sg, project 1100136). This project is supported by A*STAR under its IAF-ICP funding (Award Number I1701E0011).

## 9. Author contributions

OVM conceived the project, closely supervised the procedures, analyzed the data and wrote the manuscript; MAP and SNAA conducted the research on bacterial isolates; RRXL performed metagenomic sequencing of the clinical samples; CJH and SH planned the dietary intervention and secured the funding. All authors affirm that authorship is merited based on the International Committee of Medical Journal Editors authorship criteria.

